# Açai (*Euterpe precatoria* Mart. Arecaceae) from the Northern Region of Bolivia is not contaminated with *Trypanosoma cruzi*

**DOI:** 10.1101/2022.08.15.503748

**Authors:** OM Rollano-Peñaloza, N Roque-Marca, M Peñarrieta, P Mollinedo, C Rodriguez, D.M. Larrea-Alcázar

## Abstract

Chagas disease is a very important public health problem in America. It is caused by the protozoan parasite *Trypanosoma cruzi* and transmitted by vectors such as Triatomine insects. However, oral transmission is generating more *T. cruzi* infections than vectorial transmission in Brazilian Amazonic regions, probably due to the increased consumption of tropical fruits such as açai. Açai palms have become very popular due to its good nutritional properties. Açai fruits have different sources depending of their geographical origin. Açai palms (*Euterpe oleracea*) in Brazil are cultivated, while in Bolivia grow in wild populations and belongs to a different species, the solitary açai (*Euterpe precatoria*). Only açai from Brazil has been involved in *T. cruzi* oral transmission, while Bolivian açai has been regarded as disease free. In order to verify the absence of *T. cruzi* on açai products from Bolivia, we developed a method to detect *T. cruzi* DNA by real-Time PCR with internal controls for solitary açai. In this study we show that açai good manufacturing process did not interfere with the detection of açai or *T. cruzi* DNA. Finally, we report that freshly collected açai fruits and açai frozen pulps from Bolivia were not contaminated with *T. cruzi*.

## Introduction

The Chagas disease (also known as American Trypanosomiasis) is a parasitic disease endemic of the Americas and constitutes one of the most important public health problems in the continent and Amazon regions, causing disability in infected people and killing more people per year than any other parasitic disease in the region. Currently, in Bolivia the Chagas disease is a health problem in the Bolivian Chaco, inter-Andean valleys and Amazon regions, with as much as 60% of the country territory being endemic for Chagas disease. The infection is caused by the protozoan parasite *Trypanosoma cruzi* which, upon entering the body, lodges in organ tissues where it reproduces, causing organ damage, mainly to heart and digestive system (esophagus and colon).

Currently, several forms of *T. cruzi* transmission are known, three (e.g., vectorial, congenital, and blood transfusion) are recognized as the most common and widespread, while oral transmission has been traditionally considered rather rare and scarce. Chagas vectorial transmission is caused by the bite of an infected Triatomine insect vector of the Reduviidae family (Hemiptera) commonly called vinchuca or “timbuco”, an insect that is distributed from South America to Mexico. This mode of transmission can go unnoticed for years, even decades (15-20 years) before showing any symptoms and is thus known as a silent disease.

*Trypanosoma cruzi* oral transmission is caused by consumption of food contaminated with Triatomine feces or crushed Triatomines, ingestion of meat from infected animals not properly cooked, or food contaminated by secretions from infected marsupials (*Didelphis marsupialis*) or domestic animals. After an oral infection, the incubation period ranges between 3 and 22 days, producing fever, myalgias, vomiting, and other symptoms that may persist for four to eight weeks and might lead to other complications.

In recent years, there have been a significant increase of oral transmission of *T. cruzi* by the consumption of infected tropical fruits, such as açaí (specifically of pulp obtained from *Euterpe oleracea* Mart. fruits). Studies about the probable route of infection have shown that nowadays most infections are through oral transmission (38.3%) compared to vectorial transmission (35.4%) in Brazil. It is noteworthy that the North region is responsible for 55,6-75,3% of all Brazilian cases of *T. cruzi* infection (Santos et al. 2020) (Gov.br, 2021).

Açaí has generated great interest internationally due to its high levels of antioxidants, amino acids, minerals such as calcium, zinc, magnesium, and iron, as well as their anti-inflammatory properties. In Bolivia, the species under exploitation is the solitary açaí (*Euterpe precatoria* Mart.) which is produced in the northern Bolivian Amazon region. Solitary açaí grows up to 20-25 m. in height and they are abundant in the wild, inside different types of Amazon forests from *terra firme* mature forests to seasonally flooded forests. Solitary açaí is exploited by local producer associations, consolidating itself as an important economic alternative for the small farmer families of several communities of Pando, which have formed productive associations to market this product under principles of food safety.

Oral transmitted *T. cruzi* infection through consumption of açaí juice has been detected in several locations of Brazil but has not been reported in Bolivia yet. There are reports of oral transmitted infections in the Brazilian municipalities of Coari (Finamore-Araujo et al. 2021), Manaos (Santana et al. 2019), Amazonas State, Pará State (de Mattos et al. 2017) and Amapá State (Rodrigues 2015). However, most of these municipalities are still far from the Bolivian border, with few reports (less than 10 cases) in the neighbor states of Acre and Rondonia (Shikanai-Yasuda and Carvalho 2012).

According to the guidance for prevention measures of foodborne Chagas disease, qPCR is the best detection method of *T. cruzi* in contaminated foods (de Oliveira et al. 2019). Therefore, we developed a Real-Time PCR protocol to evaluate the presence of *T. cruzi* in raw berries and açai frozen pulps from the Northern region of Bolivia. Results indicate that *T. cruzi* is not present in açai samples from these regions.

## Materials and Methods

### Biological Materials

Solitary açai (*Euterpe precatoria*) fruits were harvested during 2020 and 2021 dry seasons (April-August) from wild palms that inhabit the *terra firme* and flooded forest at the northern region of Bolivia, in the Department of Pando (Fig. 1).

**Figure 1.**
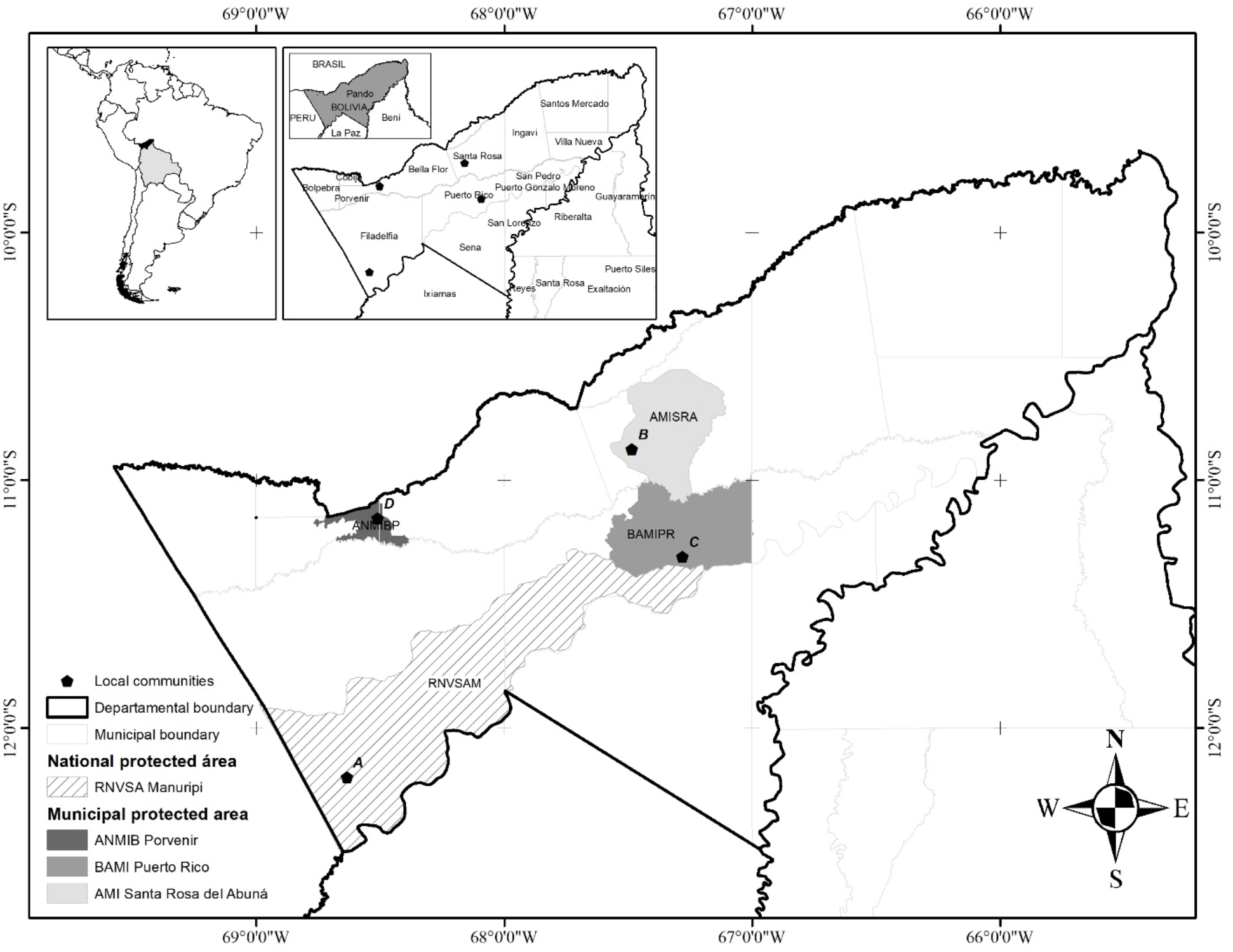
Map of Pando, Bolivia, showing açai sampling sites of four communities with productive initiatives dedicated to açai pulp commercialization. A: Villa Florida, Ventana Amazonica, B: 1ro de Mayo, ASICOPTA, C: Jericó, AFIPA-CJ and D: Trinchera, ARPTFAT. RNVSA-M: Reserva Nacional de Vida Silvestre Amazónica Manuripi, AMI-SRA: Área Modelo de Manejo Integral del Bosque Santa Rosa del Abuná, BAMI-PR: Bosque Amazónico de Manejo Integral Puerto Rico, ANMIB-P: Área Natural de Manejo Integrado del Bosque de Porvenir. The complete list of communities and productive initiatives can be found on Suppl. Table 1.

Frozen açai pulps were prepared from the 2020 harvest from 4 different communities that possess their own food processing plants (Trinchera, 1ro de Mayo, Jerico and Villa Florida) (Fig. 1, Suppl. Table 1). Açai berries for one sample (Cobija) were collected from Villa Florida community and processed at the Cobija city food processing plant. Samples were collected from each food processing plant and they were transported to the laboratory of UMSA in La Paz, maintaining a cold chain. Raw fruits were collected from the 2021 harvest from 3 communities (1ro de Mayo, Trinchera and Jerico) and immediately transported to La Paz (Fig. 1, Suppl. Table 1).

The Tulahuen strain *of Trypanosoma cruzi* (TcII) was kindly provided by Dr. Celeste Rodriguez (Immunoparasitology Unit, Department of Pathology, Faculty of Medicine, UMSA) *Trypanosoma cruzi* parasites were obtained from sediments of an *in vitro* culture with a cryoprotectant solution. *T. cruzi* parasites were maintained in VERO cells grown in Dulbecco’s Modified Eagle’s culture medium (DMEM, Merck, USA). Cryopreservation was performed as described by Martin et al. (2014).

### Açai sample collection

Açai fruits and seeds were collected before and after the washing process at the different production factories of each productive initiative. Açai fruits on arrival at the lab were placed in a beaker with sterile miliQ water in a 2:1 ratio and mixed. The residual water was filtered on a 47mm PDVF membrane with a 0.22 µM pore size. Immediately after filtration, the filters were collected for DNA extraction.

Frozen açai fruit pulps were stored at -20°C after its production in the productive initiative factories. One kilogram of frozen açai pulp was shipped, keeping the cold chain. Upon arrival at the laboratory they were thawed and 3 samples from the top, middle and bottom of the bag were collected. Samples were homogenized in a plate and 100ul of the mixed sample were collected for DNA extraction.

### DNA extraction and quantification

DNA extractions were done with the Purelink Genomic DNA kit (Thermo, USA) with the following modifications during sample lysis: Açai fruits DNA extraction was performed from PDVF filters containing wash-residues. PDVF filters were soaked in 600 ul lysis buffer, homogenized by inversion 10 times and incubated at 55°C for 1 hour. Açai pulp DNA extraction was performed from homogenized frozen pulps incubated at 55°C for 1 hour without using liquid nitrogen. Cryopreserved *T. cruzi* sediments were resuspended in 200 ul lysis buffer and incubated at 55°C for 1 hour. The rest of the DNA extraction procedures were done as described by the manufacturers instructions. DNA was quantified by fluorometry using a Qubit 2.0 Fluorometer (Thermo Scientific, USA).

### Molecular detection of Açai and T. cruzi by Real-time PCR

Specific primers for Açai and *T. cruzi* were selected from previous studies (Table 1). Specificity was confirmed by BLAST and absence of primer-dimers was verified by agarose gels. Real-time PCR was performed with Fast Sybr Green Master Mix kit (Thermo, USA) supplemented with 0.25 µM of each primer with the following conditions: 1 cycle of: 95° C, 20 s; 30 cycles of: (95°C, 3s; 60°C, 30s) with a reaction volume of 10 μl. The limit of detection for *T. cruzi* was performed with a calibrated standard curve with the most sensitive gene (Tc_kDNA) with 5 dilutions of factor 10. Absence of *T. cruzi* cross-amplification with açaí DNA was verified by inoculating *T. cruzi* DNA in the PCR reaction with açaí primers. A similar assay was performed to verify that *T. cruzi* primers did not generate amplicons with Açai DNA.

**Table 1.**
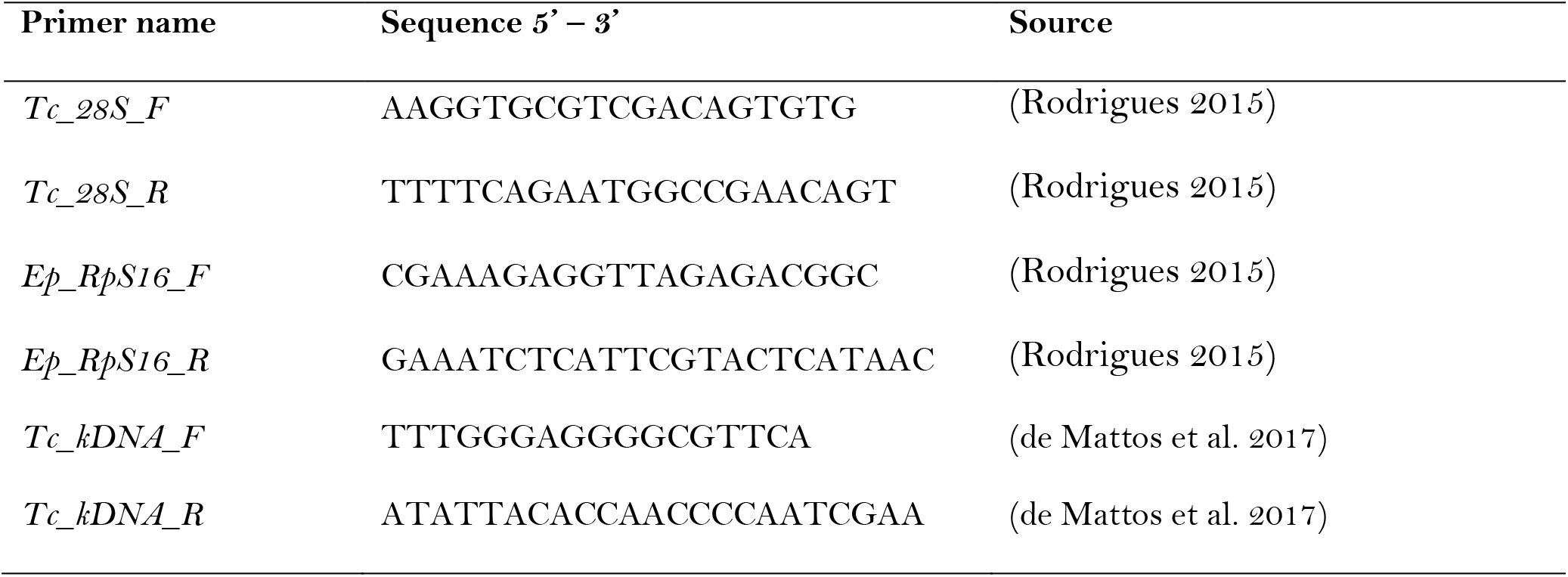
Açai and *Trypanosoma cruzi* primers utilized in this study

### Simulation of T. cruzi infection of Açai products

To simulate *T. cruzi* infection of açai products, 1ml of açaí pulp was inoculated with 10μl of purified *T. cruzi* DNA. Real-Time PCR was performed as described above.

## Results

### Validation of the molecular detection method of T. cruzi on Açai products

We first verified that our selected oligonucleotide primers for the Real-time PCR were working successfully to detect *T. cruzi* DNA with the cryopreserved parasites grown in VERO cells (Fig. 2, *T. cruzi* control). Next, we verified that our primers selected to detect Bolivian Açai (*E. precatoria*) were working with DNA extracted from frozen Açai pulps from two different productive initiatives (Trinchera y Villa Florida). We observed successful amplification of the Açai gene (Ep_16S) in those samples (Fig. 2). Finally, in order to verify that we could detect *T. cruzi* DNA in açai samples, we spiked açai frozen pulps of the same previous productive initiatives with *T. cruzi* DNA. Samples spiked with *T. cruzi* DNA shown that it was possible to detect *T. cruzi* in açai pulps with our proposed molecular method (Fig. 2).

**Fig 2.**
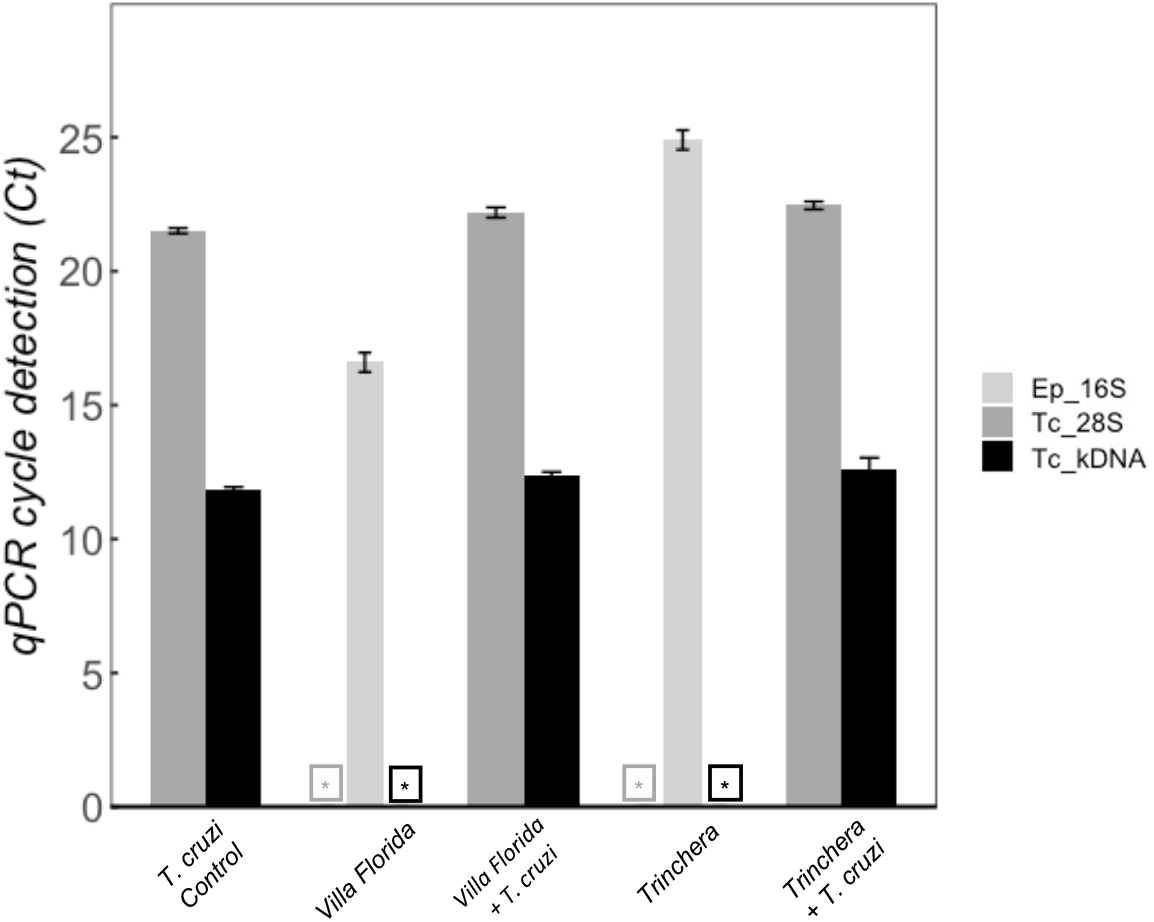
Validation of the molecular detection protocol. Açai frozen pulps were spiked with *T. cruzi* DNA. \**T. cruzi* DNA was not detected by qPCR on fresh açai frozen pulps. Tc: *T. cruzi*, Ep: *E. precatoria*

### Açai products from the Northern region of Bolivia were not contaminated with T. cruzi

After the *T. cruzi* molecular detection method with açai internal controls was validated we proceed to evaluate the presence of *T. cruzi* in frozen açai pulps from four productive initiatives (located in Trinchera, Villa Florida, 1ro de Mayo and Jerico) of the northern region of Bolivia. None of the frozen açai pulps analyzed showed presence of *T. cruzi* DNA (Fig. 3). Absence of PCR inhibitory contaminants was confirmed by the correct amplification of the açai gene *Ep_RpS16* in all samples (Fig. 3).

**Fig. 3.**
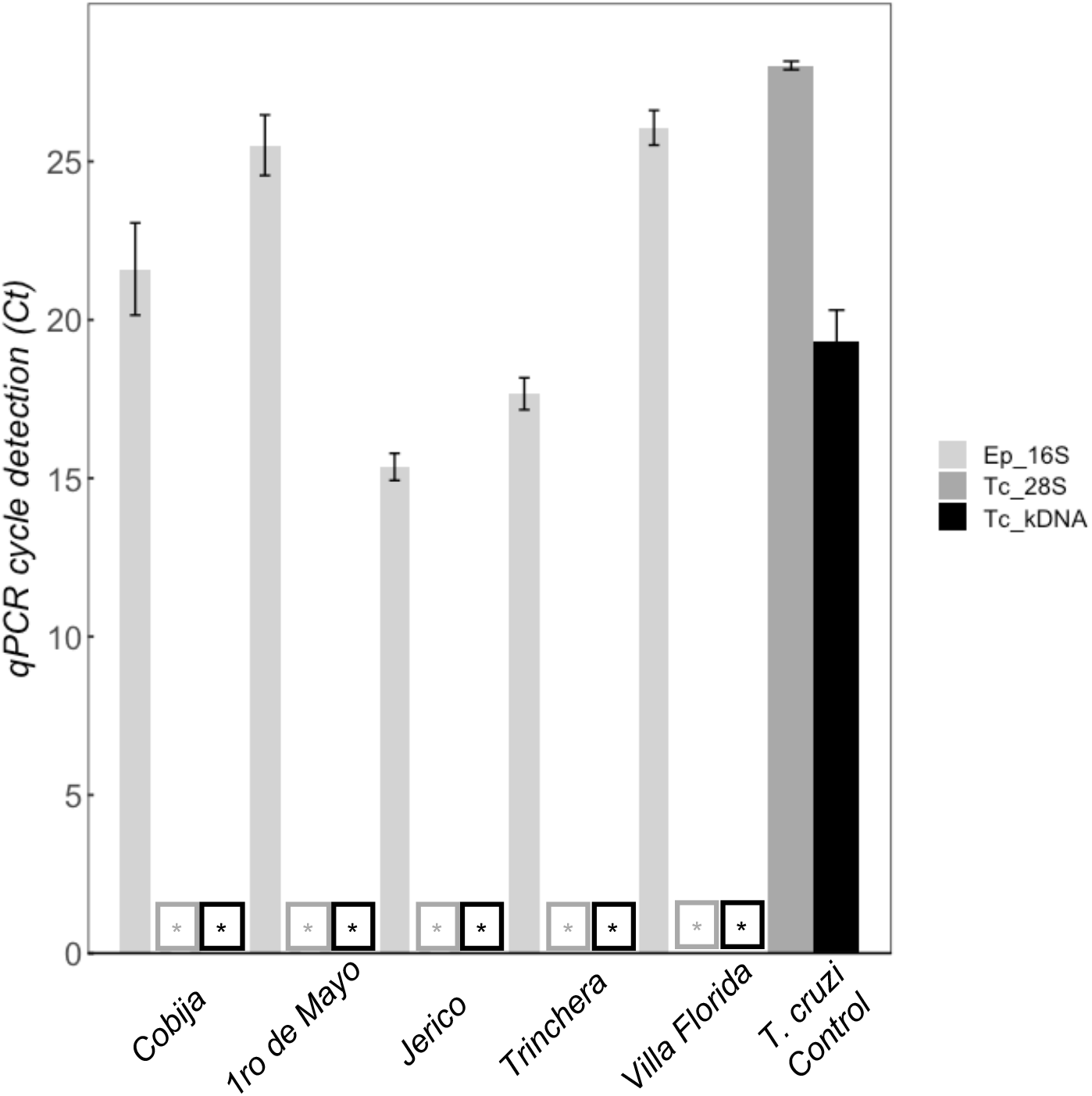
*Açai frozen pulps from Northern Bolivia* productive initiatives *do not contain Trypanosoma cruzi DNA*. \**T. cruzi* DNA was not detected by qPCR.

Good manufacturing practices for açai berries food-processing are berry washing, seed removal, seed-free pulp heat-shock, pulp freezing and frozen storage. Any of these practices might remove *T. cruzi* parasites and its genetic material, impairing the molecular detection in açai frozen pulps. We have considered that pulp heat-shock might reduce the detectability of *T. cruzi* DNA. However, it did not affect the procedure for detection, on the contrary we observed an improved detectability when açai pulps were subjected to heat treatments (data not shown). Finally, we evaluated the presence of *T. cruzi* in raw açai berries before and after the washing process from three communities that were previously analyzed as frozen pulps (1ro de Mayo, Jerico and Trinchera). We verified that raw açai fruits did not contain DNA from *T. cruzi* (Fig. 4). DNA from Açai used as an internal control was amplified in all samples.

**Fig. 4.**
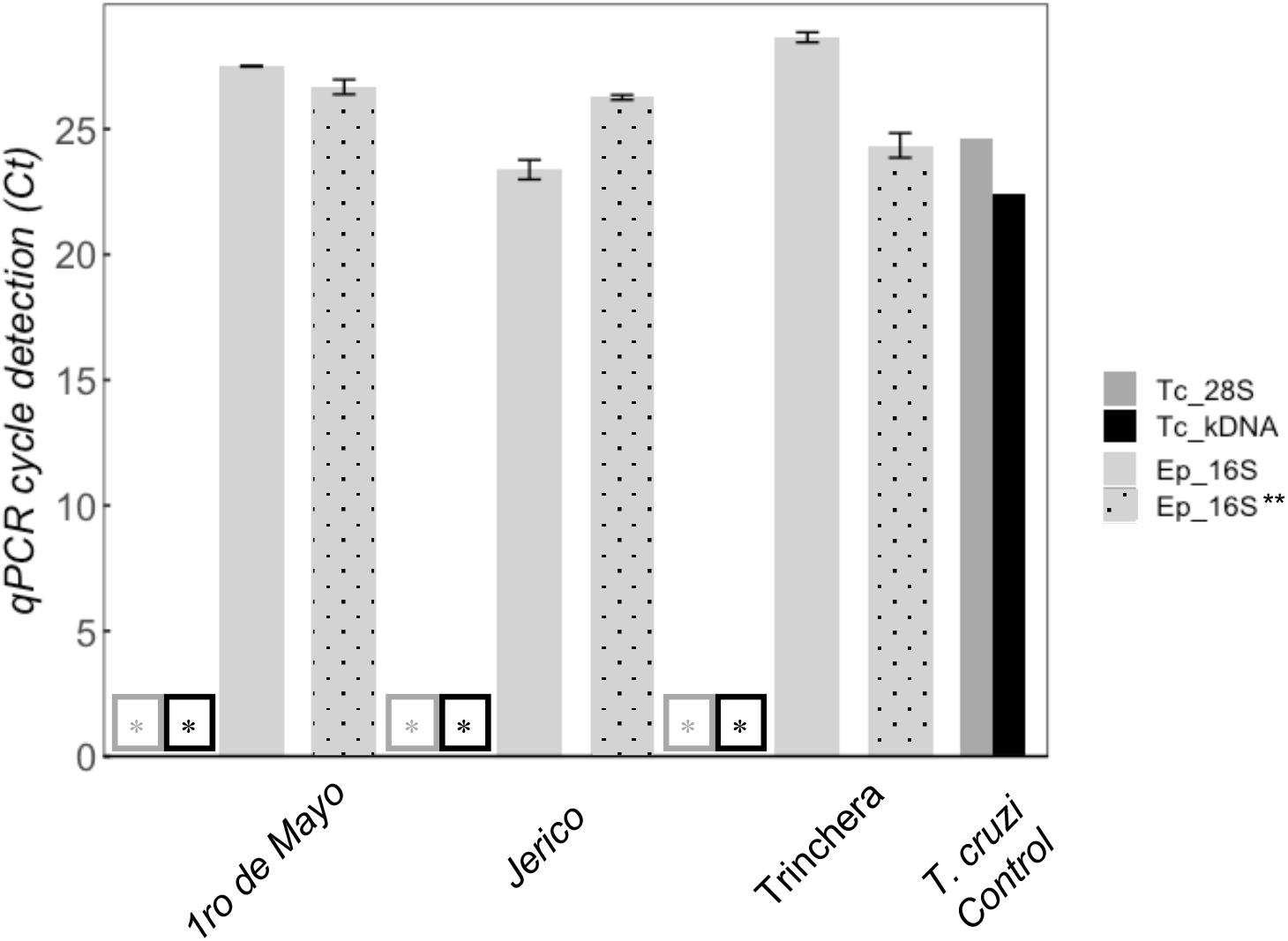
Raw açai berries from Northern Bolivia productive initiatives are not contaminated with Trypanosoma cruzi parasites. \**T. cruzi* DNA was not detected by two pairs of primers. ** Same samples were analyzed after washing in the food processing plant with similar results, only positive control is shown.

## Discussions

*Trypanosoma cruzi* oral transmission has been reported in several South American countries over consumption of tropical fruits. However, oral transmission of *T. cruzi* by consumption of açai (*E. oleracea*) has only been detected in Brazil (Velásquez-Ortiz and Ramírez 2020). *E. oleracea* grows mostly at Pará State in the Brazilian Northern region which is endemic of Chagas disease (Dias et al. 2008). This coincides with the high incidence of *E. oleracea* fruits infected with *T. cruzi*.

In Bolivia, *E. precatoria* grows in regions far away from Chagas disease endemic regions which are located in the center and south of the country (Cochabamba and Tarija valleys) (de Araújo-Jorge and Medrano-Mercado 2009).

The regions where *E. oleracea* grows in Bolivia are still relatively secluded and isolated from commercial interchange and thus the probability of transporting disease vectors inside the communities is low. Furthermore, most of the Brazilian municipalities with reported oral transmission of Chagas disease by açai consumption are still far from the Bolivian border. The closest reports are from Feijó in the Acre State with less than 10 cases (Shikanai-Yasuda and Carvalho 2012).

Heat treatments (blanching or juice pasteurization) reduces considerably the virulence of *T. cruzi* on açai fruits (Barbosa et al. 2016; de Oliveira et al. 2019). Fortunately, we have observed that heat-shock of açai pulp spiked with *T. cruzi* DNA did improve its detectability and should not be considered a burden during molecular detection.

Animal feces are a source of *T. cruzi* parasites that infects fruits involved in orally-transmitted Chagas disease (Shikanai-Yasuda and Carvalho 2012). Therefore, we assume that certain animal species might be the reason that trigger orally-transmitted Chagas disease outbreaks in localized regions and might be the reason why we do not find oral Chagas outbreaks caused by açai fruits in Bolivia yet.

Generalist animals that can thrive in a wide variety of environmental conditions can spread diseases along their path while escaping from deforestation (Dobson et al. 2020). An example of these are common opossums (*Didelphis marsupialis*) which are known to be *T. cruzi* reservoirs in Brazil but have not been recorded during infections in Bolivia (Erazo et al. 2019; Velásquez-Ortiz and Ramírez 2020). Thus, deforestation of Brazilian forest might push infected animals towards Bolivian conserved forests and generate new orally-transmitted Chagas disease outbreaks. To prevent such scenarios, our recommendation is to implement a molecular surveillance system such as Real-time PCR to detect *T. cruzi* in Bolivian tropical fruits such as açai (*E. precatoria*).

In short, our results show that raw açai fruits and açai frozen pulps from Bolivia are not contaminated with *T. cruzi*. These data can be a baseline to design a monitoring system focused on local communities and productive initiatives related to the productive chain of açai in northern Bolivia. Our results reinforce the importance of the Good Manufacturing Practices (GMP), which have to guide any biodiversity-based productive initiative linked to human health.

## Acknowledgements

This project was supported by World Wildlife Fund (WWF) through a fund provided by the Federal Ministry for Economic Cooperation and Development (BMZ) of Germany, and Andes Amazon Fund (AAF). We are grateful to the Immunoparasitology Unit of the Faculty of Medicine for the provision of cryopreserved *T. cruzi* st. *Tulahuen* (TcII).

