## Supplementary Table 1 for "Açai (*Euterpe precatoria* Mart. Arecaceae) from the Northern Region of Bolivia is not contaminated with *Trypanosoma cruzi*"

Supplementary Table 1. Açai sampling sites of four communities with productive initiatives dedicated to açai pulp commercialization in Pando Department, Bolivia.

| * | *Community name* | *Productive initiative* | *Abbrev.* | *National protected Area* | *Abbrev.* | *Frozen pulp qPCR analysis* | *Raw fruits qPCR analysis* |
|---|---|---|---|---|---|---|---|
| A | Villa Florida | Ventana Amazónica** | --- | Reserva Nacional de Vida Silvestre Amazónica Manuripi | RNVSA-M | Yes** | --- |
| B | 1ro de Mayo | Asociación Integral de Cosechadores, Productores y Transformadores de Frutos del Abuná | ASICOPTA | Área Modelo de Manejo Integral del Bosque Santa Rosa del Abuná | AMI-SRA | Yes | Yes |

| | | | | | | | |
|---|---|---|---|---|---|---|---|
| C | Jericó | Asociación Forestal Integral de Productores Agropecuarios de la Comunidad de Jericó | AFIPA-CJ | Bosque Amazónico de Manejo Integral Puerto Rico | BAMI-PR | Yes | Yes |
| D | Trinchera | Asociación de Recolectores, Productores y Transformadores de Frutos Amazónicos de Trinchera | ARPTFAT | Área Natural de Manejo Integrado del Bosque de Porvenir | ANMIB-P | Yes | Yes |

** Geographical position can be observed in Pando Department map, Bolivia (Fig. 1).*

*** Productive initiative Ventana Amazónica has two processing plants, one located at Villa Florida community and one at Cobija, the Capital of Pando department.*
